# Integrated biorefinery from *Arthrospira platensis* biomass as feedstock for bioethanol and lactic acid production

**DOI:** 10.1101/715672

**Authors:** Diego A. Esquivel-Hernández, Anna Pennacchio, Roberto Parra Saldivar, Vincenza Faraco

**Author notes:** Corresponding author: Vincenza Faraco.

## Abstract

An integrated biorefinery for ethanol and lactic acid production from the biomass of cyanobacterium *Arthrospira platensis* was investigated. Different pretreatments consisting of supercritical fluid extraction (SFE) and microwave assisted extraction (MAE) with non-polar (MAE-NPS) and polar solvents (MAE-PS) were tested on cyanobacterial biomass to obtain bioactive metabolites and the resulting residual biomass was used as a substrate for fermentation with *Saccharomyces cerevisiae* LPB-287 and *Lactobacillus acidophilus* ATCC 43121 to produce ethanol and lactic acid, respectively. The maximum concentrations achieved in our processes were 3.02±0.07 g/L of ethanol by the MAE-NPS process at 120 rpm 30 °C, and 9.67±0.05 g/L of lactic acid by the SFE process at 120 rpm 37 °C. Our results suggest that the proposed approach can be successfully applied in bioactive metabolites extraction and subsequently in the production of Ethanol and Lactic acid from *A. platensis* depleted biomass.

## Introduction

The cyanobacterium *Arthrospira platensis* is an outstanding sunlight-driven cell factory that has the ability to convert carbon dioxide into high added value metabolites (Singh et al., 2017). This photosynthetic cyanobacterium have received much attention for high value metabolites such as α-tocopherol, β-carotene, lutein and γ-linolenic acid (Esquivel-Hernández et al., 2016; Sosa-Hernández et al., 2019) and bioactive effects such as antiproliferative, antitumor, antifungal, antibacterial, antimalarial, antiviral, antimycotic, cytotoxicity, multi-drug resistance reversers and immunosuppressive agents (Dixit & Suseela, 2013; Ediriweera et al., 2017; Kumari et al., 2011).

*A. platensis* contains approximately 65% of proteins, 20% of carbohydrates, 7% of minerals and 5% of lipids, (Campanella et al., 1999). After extraction of the high value metabolites, the resulting residual biomass, containing polysaccharides, monosaccharides and proteins (Esquivel-Hernández et al., 2017), can be a suitable feedstock for production of ethanol and lactic acid that can be used as biofuels and raw material of biopolymers in the frame of biorefinery concept and circular economy (Esquivel-Hernández et al., 2016).

A biorefinery is an industrial plant, in which all the components of biomass are converted into a range of biochemicals, materials, and energy products, and byproducts of a process are used as raw materials for another production process, aiming at a zero-waste biorefinery as pillar of circular economy (Liguori et al., 2013; Liguori & Faraco, 2016). This approach can be applied to improve the economic balance of biofuel production from cyanobacteria, through co-production of high-value metabolites from the same microorganisms, generating an integrated biorefinery for sustainable production of biofuels while exploiting other traditional cyanobacterial applications.

The lack of lignin in the cell wall of *A. platensis* entails milder pre-treatments for releasing the fermentable sugars than lignocellulosic biomasses, having therefore the potential to overcome the problems related to pretreatments with other currently adopted feedstock (Capolupo & Faraco, 2016).

To date, few studies have been reported on bioethanol production from *A. platensis*. An interesting approach is the saccharification of the carbohydrate enriched biomass and the highest ethanol production of 2.03 of g/L was obtained from the hydrolysate of the biomass with nitric acid (0.5 N) (Markou et al., 2013). A direct conversion of *A. platensis* to ethanol mediated by the use of lysozyme and a recombinant amylase-expressing yeast strain has produced a concentration of ethanol of 6.5 g/L (Aikawa et al., 2013). The technical feasibility of the bioethanol production from *A. platensis* has been evaluated with a percentage of bioethanol among 0.85-1% (w/w) and by the measurement of the variation of ethanol percentage in the final solution according to the variation of air-drying time using three different air-drying times (1 day, 2 days and 3 days air drying) (Hossain et al., 2015).

In the case of lactic acid, to the best of our knowledge, the information of the use of *A. platensis* biomass as feedstock for production of lactic acid is scarce. Lyophilized biomass from *A. platensis* has been used as substrate for lactic acid fermentation by the probiotic bacterium *Lactobacillus plantarum* ATCC 8014. After 48 h of fermentation the lactic acid concentration reached 3.7 g L^−1^ (Niccolai et al., 2019).

Based on this, the aim of this work was to analyze the strains *Saccharomyces cerevisiae* LPB-287 (Liguori et al., 2015a) and *Lactobacillus acidophilus* ATCC 43121 (Liguori et al., 2015b) for their ability to grow on the *A. platensis* residual biomass after high value metabolites extraction by either supercritical fluid extraction (SFE) or microwave-assisted extraction (MAE) and produce ethanol and lactic acid, respectively.

## Materials and methods

### Materials

Ammonium sulfate (NH_4_SO_4_), potassium phosphate dibasic (K_2_HPO_4_), potassium phosphate monobasic (KH_2_PO_4_), zinc chloride (ZnCl_2_), magnesium sulfate (MgSO_4_), 3,5-Dinitrosalicylic acid (DNS), sulfuric acid (H_2_SO_4_), potassium hydroxide (KOH), glucose (C_6_H_12_O_6_), bradford reagent and yeast extract were obtained from Sigma Chemical Co., (St. Louis, MO, USA). Peptone and malt extract were obtained from BD Biosciences (Heidelberg, Germany). MRS broth from Oxoid Ltd (Basingstoke, United Kingdom).

### Cyanobacteria cultivation

The cyanobacterial strain *Arthrospira platensis* used throughout this work was cultivated as reported in (Esquivel-Hernández et al., 2017a) (Esquivel-Hernández et al., 2017b). Briefly, *A. platensis* was grown for 45 days in open raceway ponds with modified Jourdan medium whose composition in (g/L^-1^) is as follows: NaHCO3 (5.88), Na3PO4 (0.16), NaNO3 (0.92), MgSO4·7H2O (7.07), FeSO4 (0.004) and NaCl (2). The geographical location of the ponds was 20″1410” N,103″3510” W. The biomass was harvested with a mesh, air dried to 20% moisture and stored under dry and dark conditions until analysis. The complete frame of integrated biorefinery appears on Figure 1.

**Fig. 1.**
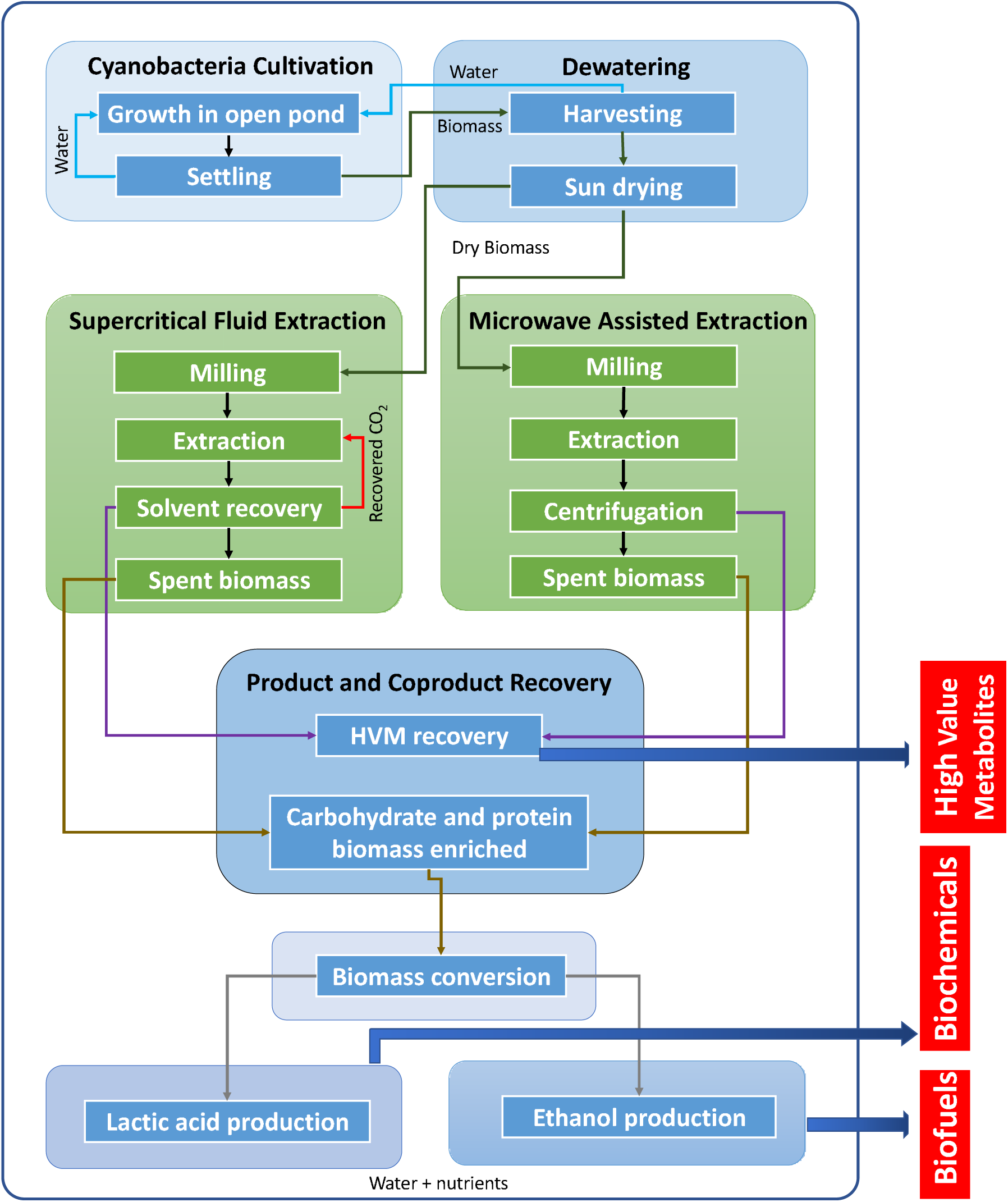
Cyanobacteria biomass and by-products conversion into biofuels, biochemicals and high value metabolites through green extraction technologies and fermentation processes in the frame of an integrated biorefinery.

### Extraction of high value metabolites from cyanobacteria by supercritical fluid extraction (SFE)

Supercritical fluid extraction (SFE) of high value metabolites from the *A. platensis* cells was performed according to our previous work (Esquivel-Hernández et al., 2017a). Briefly, all extractions were carried out using a 100 mL extraction cell (Thar SFC SFE 100, Waters Corp., Milford, MA, USA). All extractions were performed at a 25 g/min CO_2_ flow and ethanol as co-solvent with a Plackett–Burman design reported in Table 1(a). A Plackett–Burman design was used for the six experimental factors generating 12 experimental conditions tested with triplicates carried out in randomized run order. In this design, all six factors were tested at two different experimental levels: Co-solvent (CX) (g/min 4, 11); pressure (P) (bar 150, 450); static extraction (SX) (min 5, 15); dynamic extraction (DX) (min 25, 55); temperature (T) (°C 40, 60); and, dispersant (Di) (g 0, 35).

The results of each response variable and their statistical significance were analyzed with ANOVA (p > 0.05), using the statistical package Minitab 16 (State College, PA, USA). The cyanobacterial residual biomass was dried at 60 °C in an oven in order to eliminate impurities that were generated during the SFE process before fermentation process.

**Table 1.** Experimental designs for green extraction processes. a) Supercritical Fluid Extraction (SF) b) Microwave assisted Extraction MAE with polar (MP) and non-polar solvents (MN).

| Run | CS<br>(g/min) | P<br>(bar) | SE<br>(min) | DE<br>(min) | T<br>(°C) | DS<br>(g) |
| --- | --- | --- | --- | --- | --- | --- |
| SF1 | 4 | 450 | 5 | 25 | 40 | 35 |
| SF2 | 11 | 450 | 15 | 25 | 60 | 35 |
| SF3 | 11 | 450 | 5 | 55 | 60 | 0 |
| SF4 | 4 | 150 | 5 | 55 | 60 | 35 |
| SF5 | 11 | 150 | 15 | 55 | 40 | 35 |
| SF6 | 4 | 150 | 15 | 55 | 60 | 0 |
| SF7 | 4 | 150 | 5 | 25 | 40 | 0 |
| SF8 | 11 | 450 | 5 | 55 | 40 | 0 |
| SF9 | 11 | 150 | 15 | 25 | 40 | 0 |
| SF10 | 4 | 450 | 15 | 55 | 40 | 35 |
| SF11 | 11 | 150 | 5 | 25 | 60 | 35 |
| SF12 | 4 | 450 | 15 | 25 | 60 | 0 |

Co-solvent (CX), pressure (P), static extraction (SX), dynamic extraction (DX), temperature (T), dispersant (Di). b) Microwave assisted Extraction MAE with polar (MP) and non-polar solvents (MN).
| Run | Run | S<sup>1</sup><br>(v/v) | t<br>(min) | T<br>(°C) |
| --- | --- | --- | --- | --- |
| MP1 | MN1 | 0.25 | 15 | 40 |
| MP2 | MN2 | 0.25 | 15 | 60 |
| MP3 | MN3 | 0.25 | 55 | 40 |
| MP4 | MN4 | 0.25 | 55 | 60 |
| MP5 | MN5 | 0.81 | 15 | 40 |
| MP6 | MN6 | 0.81 | 15 | 60 |
| MP7 | MN7 | 0.81 | 55 | 40 |
| MP8 | MN8 | 0.81 | 55 | 60 |

<sup>1</sup> Solvent ratio refers to ammonium acetate 10 mM and ethanol (v/v) for MP and limonene and ethyl acetate for MN, Time (Ti), and Temperature (Te).

### Extraction of high value metabolites from cyanobacteria by microwave-assisted extraction (MAE)

Microwave-assisted extraction (MAE) of high value metabolites from the *A. platensis* cells was performed according to our previous work (Esquivel-Hernández et al., 2017b). Briefly, MAE experiments were carried out in a microwave-assisted extraction equipment MARS 5 (CEM Corporation, Matthews, USA) with a 100 mL extraction vessel Green-Chem. The extraction vessel was built with PFA Teflon. All extractions were done at 400W power and 1 bar. For each experiment, the extraction vessel was filled with a 0.14 ratio of dry weight biomass/solvent (w/v). The solvent ratio was defined as a v/v ratio (v/v; 0.25, 0.81) of the combination of two types of solvents, polar solvents MP (Ammonium acetate 10 mM and ethanol) and non-polar solvents MN i.e., Limonene (1-methyl-4-(1-methyle thenyl)-cyclohexene) and ethyl acetate with a 2^k^ design based on three experimental factors generating 8 experimental conditions (1-8) tested with triplicates carried out in randomized run order with the experimental matrix design reported in Table 1(b). The cyanobacterial residual biomass was dried at 60 °C in an oven in order to eliminate impurities that were generated during the MAE process before fermentation process.

### Analysis of sugar content of residual cyanobacterial biomass

In order to determine the cyanobacterial biomass composition, the samples of SF, MN and MP pretreatments with highest yields for ethanol and lactic acid, respectively (Table 1; Fig 3, 7) were selected and subjected to acid hydrolysis. 5 g of lyophilized cyanobacterial depleted biomass were mixed with sulfuric acid to reach at a final acid concentration of 0.1% (v/v). The resulting slurries were then autoclaved at 121 °C for 20 min. After hydrolysis, the samples were cooled to room temperature, centrifuged at 4 °C and 9000*g* for 20 min and the supernatant containing the released sugars was collected as acidic hydrolysate, the sugar content and composition of which was measured by HPLC method as described in paragraph Analytical Methods.

### *S. cerevisiae* inoculum and medium preparation for fermentation and ethanol production

The yeast strain adopted for ethanol fermentation was *S. cerevisiae LPB-287* selected in our previous work (Liguori et al., 2015a). The yeast was inoculated in three 250 mL flasks containing 100 mL of YM broth whose composition in (g/L^-1^) is as follows: glucose (10), peptone (5), malt extract (3), yeast extract (3). The medium was autoclaved before using for fermentation. The cultivation was carried out at 120 rpm for 24 h and 30 °C. After 24 h, cells were collected by centrifugation at 10,000 rpm for 15 min and washed three times in sterilized water. The medium for the yeast fermentation and ethanol production was prepared starting from that previously reported for the ethanol production from *A. platensis* (Markou et al., 2013) with slight changes. Its composition in (g/L^-1^) was as follows: NH_4_SO_4_ (2), K_2_HPO_4_ (1), KH_2_PO_4_ (1), ZnSO_4_ (0.2), MgSO_4_ (0.2) and yeast extract (2). Fermentations were carried out with the strain S. cerevisiae *LPB-287* in 250 mL Erlenmeyer flasks. 10 g/L concentration of cyanobacterial biomass was prepared (for all samples from SFE and MAE) and added to 50 mL of sterilized fermentation medium. 5 mL of yeast culture was harvested in the exponential phase and used to inoculate each fermentation medium aseptically. The flasks were incubated at 30 °C under 120 rpm. The fermentation continued for 81 h and samples for analysis were taken every 24 h.

### *L. acidophilus* inoculum and medium preparation for fermentation and lactic acid production

The *lactobacillus* strain adopted for lactic acid production was *L. acidophilus* ATCC 43121 selected in our previous work (Liguori et al., 2015b). De Man, Rogosa and Sharpe broth (MRS) was used as the growth medium for the inoculum. The composition of MRS medium in (g/L^-1^) was as follows: glucose (20), peptone (10), meat extract (8), yeast extract (2), Triammonium citrate (2), K_2_HPO_4_ (2), CH_3_COONa3H_2_O (5), MgSO_4_*7H_2_O (0.2) and MnSO_4_*4H_2_O (0.05). The medium was autoclaved before using for fermentation. The strain was inoculated in three 250 mL flasks containing 100 mL of YM broth and cultivated by static incubation for 24 h and 37 °C. After 24 h, cells were collected by centrifugation at 10,000 rpm for 15 min and washed three times in sterilized water. The medium for the lactic acid production was prepared starting from that previously adopted for lactic acid production from *Hydrodictyon reticulatum* (Nguyen et al., 2012) with slight changes. The composition in (g/L^-1^) was as follows: Peptone (3), yeast extract (3). The medium was autoclaved before using for fermentation.

Lactic fermentations were carried out with the strain *L. acidophilus* ATCC 43121 in 250 mL Erlenmeyer flasks. 20 g/L concentration of cyanobacteria substrates were prepared (for all samples from SFE and MAE) and added to 50 mL of sterilized fermentation medium. 5 mL of lactobacillus culture was harvested in the exponential phase and used to inoculate each fermentation medium aseptically. The flasks were incubated at 37 °C under 120 rpm. The fermentation continued for 81 h and samples for analysis were taken every 24 h.

## Analytical methods

Reducing sugar concentration was analyzed using the dinitrosalicylic (DNS) method (Miller, 1959). Briefly, glucose was used as standard for this analysis. 200 μL of the samples were mixed with 800 μL of the DNS reagent. Tubes were placed in boiling water bath for 5 min, transferred to ice to rapidly cool down and then brought to room temperature by placing them in water bath at 25 °C. The absorbance was measured at 575 nm, using Novaspec II spectrophotometer.

Ethanol was determined using commercial kits (K-ETOH) from Megazyme International (Wicklow, Ireland). The calculations were done with Megazyme Mega-Calc and expressed as g ethanol/L. Lactic acid was determined using commercial kits (KDLATE) from Megazyme International (Wicklow, Ireland). The calculations were done with Megazyme Mega-Calc and expressed as g lactic acid/L. The monomeric sugars content was determined by high performance anion-exchange chromatography (HPAEC; DX-300 series chromatography system, Dionex, USA). The effluent was monitored with pulsed amperometric detection detector (PAD, Dionex, CA, USA). The 10 μl of filtered samples were injected into a CarboPac PA-1 anion-exchange column (0.4 x 250 mm, Dionex, CA, USA) that was pre-equilibrated in 18 mM KOH. Chromatographic separation of the monomeric sugars from the samples was achieved in the isocratic mode with 18 mM KOH at a flow rate of 1.0 ml/min in 20 min. The content of proteins was determined by the Bradford method with the Bradford reagent, and the OD of it was measured at 595 nm. Each measurement was repeated at least three times and averaged.

## Results and discussion

### Effect of supercritical fluid extraction (SFE) and microwave-assisted extraction (MAE) pretreatment of cyanobacterial biomass on ethanol production and consumption rate of reducing sugars

An integrated biorefinery for ethanol and lactic acid production from the biomass of cyanobacterium *Arthrospira platensis* was investigated. Different pretreatments consisting of supercritical fluid extraction (SF) and microwave assisted extraction (MAE) with non-polar (MN) and polar solvents (MP) were tested on cyanobacterial biomass to obtain bioactive metabolites and the resulting residual biomass was used as a substrate for fermentation with *Saccharomyces cerevisiae* LPB-287 and *Lactobacillus acidophilus* ATCC 43121 to produce ethanol and lactic acid, respectively.

For SF pretreatment, twelve types of samples were used as a substrate for ethanol fermentation by the yeast strain *S. cerevisiae* LPB-287 after the extraction of high value metabolites. A Plackett–Burman design was used to allow the selection of different factors of a SF process. The factors considered for this study were: pressure, temperature, co-solvent (ethanol), dispersant agent (glass pearls), and static and dynamic extraction. As shown in Fig 2, no differences were observed in the ethanol concentrations in the time for all samples, with a content of ethanol of 2.0 ± 0.08 g/L. About the ethanol yield, the highest yield (% gEthanol/gCyanobacteria) among the experiments was 0.20 Fig 3.

**Fig. 2.**
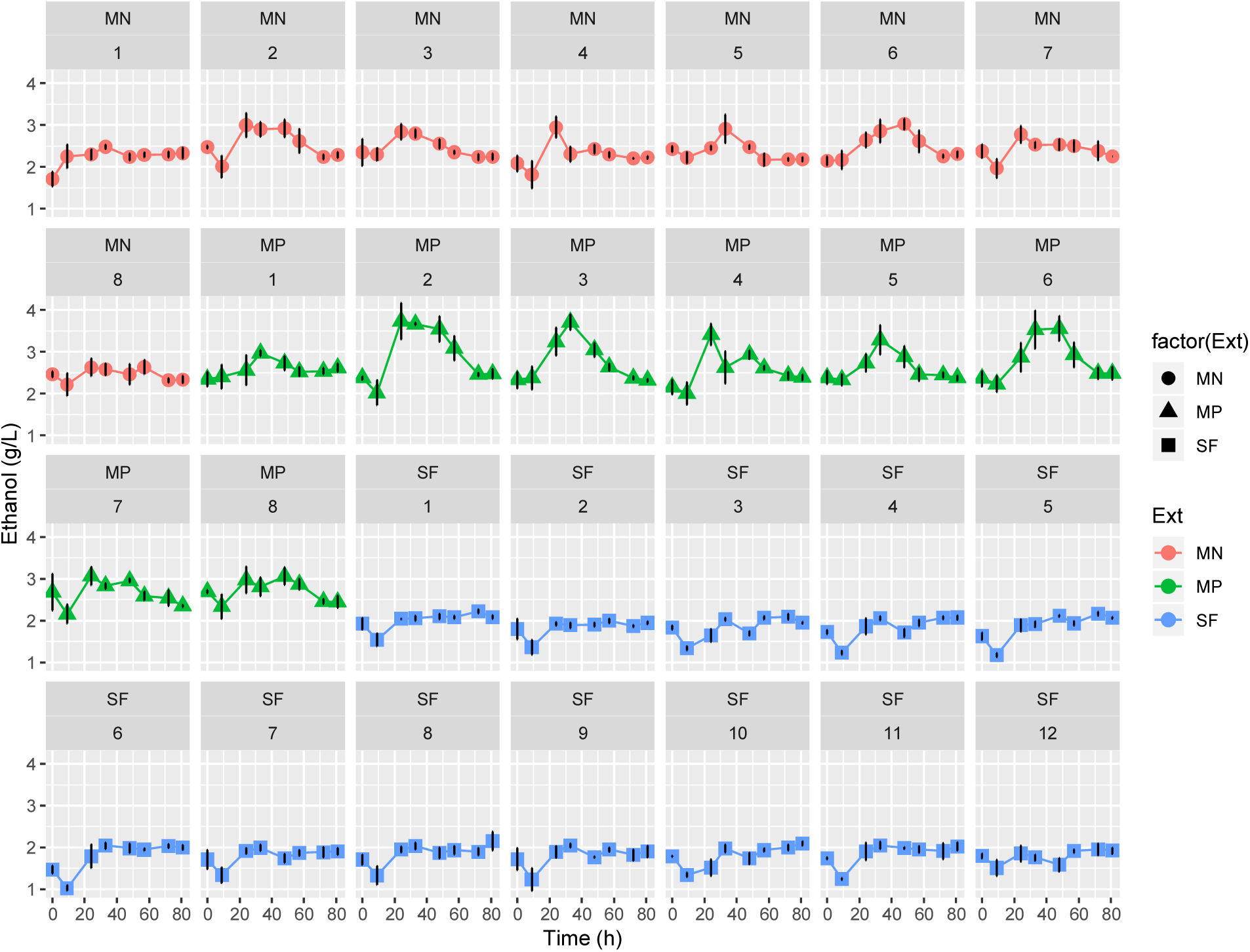
Time course of concentration of ethanol during fermentation of *Saccharomyces cerevisiae LPB-287* with depleted cyanobacteria biomass from SF, MN and MP pretreatments. *(factor(Ext) refers to the type of pretreatment respectively, n=3).

Whereas MAE pretreatment, two sets of conditions with polar (MP) and non-polar solvents (MN) were applied and the corresponding eight types of samples were adopted as substrates for ethanol fermentation by the strain *S. cerevisiae* LPB-287. As shown in Fig 2, for MN set, the highest ethanol concentrations were obtained in the experiment MN6 (3.02±0.07 g/L) and experiment MN2 (2.99±0.07 g/L). The content of ethanol is higher with MN pretreatment respect to the results with SF pretreatment. This can be due to the use of non-polar solvents for MN pretreatment that have already depleted the biomass and this residue was enriched on carbohydrates. Respecting the ethanol yield, the highest and the average yield (% gEthanol/gCyanobacteria) among the experiments was 0.30, Fig 3.

For the MP experiments, the higher ethanol concentrations were obtained at the experiment MP2 and MP3 (0.37 g/L) Fig 2. In relation to the ethanol yield, the highest value among the experiments MP2 and MP3 was 0.37 (% gEthanol/gCyanobacteria) Fig 3. These results were obtained at S=0.25, t=15 min, T=60 °C and S=0.25, t=55 min, T=40 °C conditions of pretreatment process as shown in Table 1 (b). The maximum concentrations achieved in our processes were 3.02±0.07 g/L of ethanol by the MN process at 120 rpm 30 °C. There are not studies of bioethanol production from *A. platensis* depleted biomass (with SF, MN and MP) using *Saccharomyces cerevisiae*. Furthermore, many studies have been focused on the production of ethanol from several *A. platensis* hydrolysates obtained by the use of amylases and lysozymes (Aikawa et al., 2013). Also, in this frame, there are several studies on production of ethanol using other algal cultures. It has been reported the production of bioethanol from an hydrolysate of *Chlorella* sp. biomass (1.5% of sulfuric acid at 117°C for 20 min) with a concentration of 5.62□± □0.16□g/L of ethanol by the fermentation of *S. cerevisiae* TISTR 5339 (Ngamsirisomsakul et al., 2019). A microalgae mixed culture (obtained from a freshwater area in Osku located in northwest of Iran) was pretreated with an enzymatic mixture (B-glucosidase, a-amylase and amyloglucosidase) and evaluated to produce bioethanol by fermentation with *S. cerevisiae* (ATCC 7921), the obtained results showed that the highest content of ethanol was 6.41 g/L (Shokrkar et al., 2017). Biomass from *Porphyridium cruentum* (KMMCC-1061) was subjected to an enzymatic hydrolysis with pectinase and cellulase enzymes and therefore was used as carbon source for a fermentation process with *S. cerevisiae* KCTC 7906, the results showed that the highest content of ethanol was 0.65 g/L (Kim et al., 2017).

Based on this, in comparison with *P. cruentum* cultures from Kim et al., 2017 our results are higher but in comparison with Ngamsirisomsakul et al., 2019 our results are lower. This behavior can be due to the chemistry behind the method used for pretreatment, since the use of water as solvent extraction can change the composition of monosaccharides in the depleted cyanobacterial biomass, as can be seen on Fig 4. where the control has a lower content of reducing sugars.

**Fig. 3.**
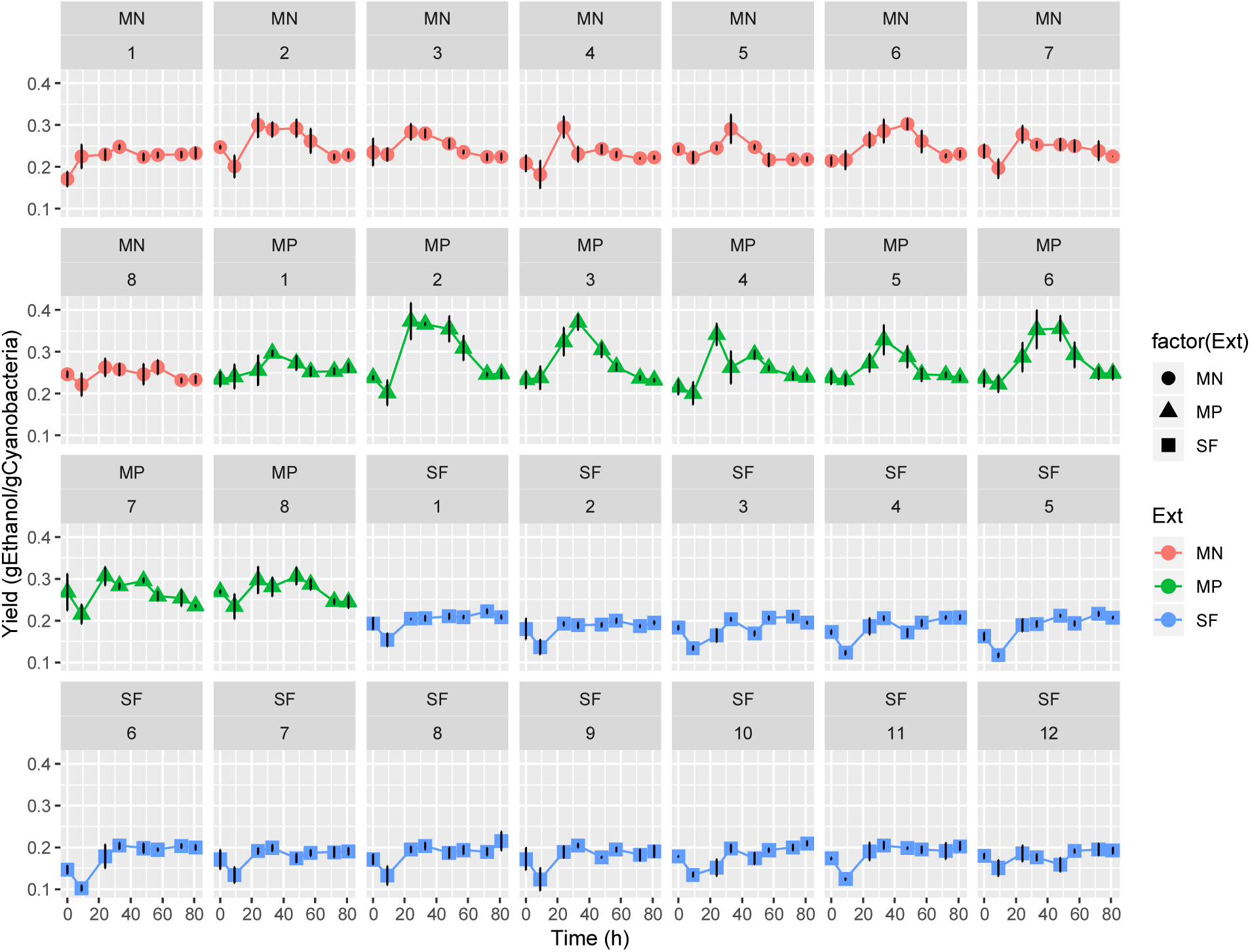
Time course of yield of ethanol during fermentation of *Saccharomyces cerevisiae LPB-287* with depleted cyanobacteria biomass from SF, MN and MP pretreatments. *(factor(Ext) refers to the type of pretreatment respectively, n=3)

**Fig. 4.**
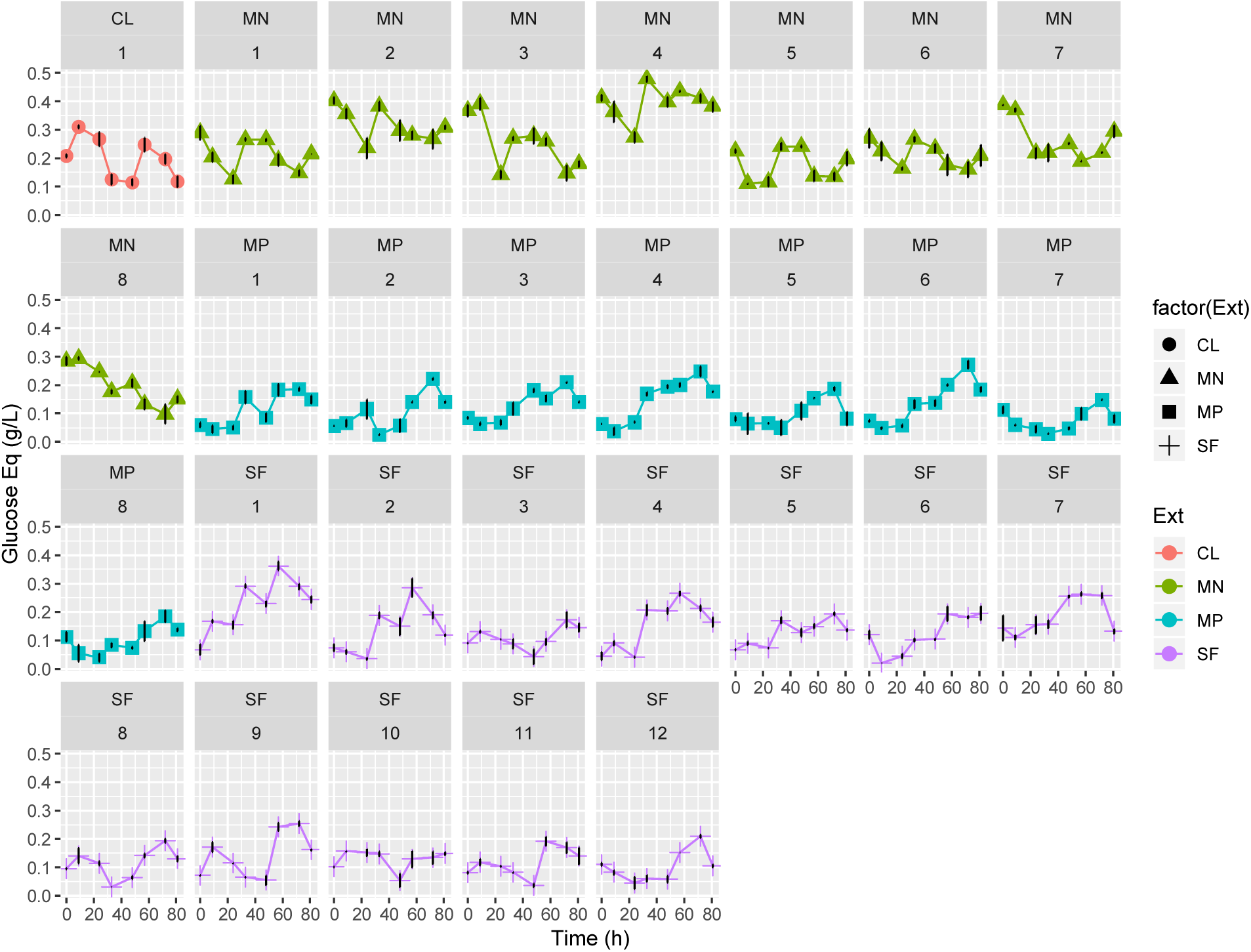
Time course of reducing sugar concentration measured by DNS method during fermentation of *Saccharomyces cerevisiae LPB-287* with cyanobacteria from SF, MN and MP pretreatments. *(factor(Ext) refers to the type of pretreatment, respectively, n=3, CL is a positive control).

In regard to the composition of monosaccharides from depleted cyanobacterial biomass, it has been reported that these microorganisms have different types of polysaccharides, peptoglycanes in the wall cell and glycogen granules in the intracellular vesicles (Van Eykelenburg, 1977). During the SFE process, the cell wall of the cyanobacteria is broken due to the high temperature and pressure required for the process (Esquivel-Hernández et al., 2017a). This leads to the release of the different polysaccharides embedded in the cell wall. It has also been reported that extracted microalgae such as *Chlorococcum* sp. improve the yield of yeast biomass by providing a sugar source for the yeast growth (Harun et al., 2010; Mondal et al., 2017).

As shown in Fig. 4, all extraction samples obtained by SF decrease in reducing sugars content when compared with control biomass. This can be due to the content of water in ethanol used for SFE processes. The highest content of reducing sugars was obtained under the experiments SF8 CS= 11 g/min, P=450 bar, T=40 °C and 1 CS= 4 g/min, P=450 bar and T=40 °C Table 1 (a).

All extraction samples obtained by MN showed a remarkable increase in reducing sugars when compared with control biomass Fig. 4 due to the extraction mixture used for MN, which comprises limonene and ethyl acetate (Esquivel-Hernández et al., 2017b). All extraction samples obtained by MP show a sharp decrease in reducing sugars when compared with control biomass Fig. 4 as long as the extraction mixture used for MAE polar extraction (MP), which comprises water (10 mM ammonium acetate) and ethanol.

For MN exists a major depletion in the reducing sugars after the first 24 hours; in this case, we can assume that hexose sugars are rapidly metabolized since *S. cerevisiae* possess carbon catabolite repression (CCR) with mixed sugars obtained from algal cultures (Kim et al., 2010). This result can be useful for the scale up of the process, since we can obtain a higher concentration of ethanol in short time. Also, it is important to distinguish that these samples exhibited a different behavior of the control sample, mainly because in this pretreatment we started at the higher level of reducing sugars, based on the polarity of the solvent used in the extraction process. The maximum content of reducing sugars was obtained under the experiments: MN2 S=0.25, t=15 min, T=60°C and MN3 S=0.25, t=55 min, T=40°C Table 1 (b).

The highest content of reducing sugars was obtained under the experiments MP8 S=0.81, t=55 min, T=60°C and MP3 S=0.25, t=55 min, T=40°C Table 1 (b). In particular we observed an increment of pentose sugars and a decrease of hexose sugars because the hexose sugars are rapidly metabolized as a result of CCR.

The main sugars in *A. platensis* biomass are glucose, xylose, mannose, fucose and rhamnose. Owing to CCR, the hexose sugars are rapidly metabolized by *S. cerevisiae*. However, because we have other kind or sugars, *S. cerevisiae* uses pentose sugars when the hexoses content has been consumed (Wang et al., 2017). Kinetics of assimilation of this mixture of sugars has cycles of 24 hours for the consumption of hexose and pentose sugars. Hexose sugars are considered to be efficiently fermented to ethanol, whereas pentose sugars were consumed after the depletion of hexose sugars (Kim et al., 2010).

Therefore, the biomass resulting from MP process is the least suitable for the ethanol production in the frame of integrated biorefinery due to low values of concentration of sugars. There is a potential for using this depleted biomass in other bioprocesses such as biogas or bioplastics production.

### Time Course of monosaccharides and protein in selected conditions of yeast fermentation

The time course of monosaccharides and protein of the most representative samples of SF, MN and MP pretreatments in the fermentation by *S. cerevisiae* was performed. For SF, the experiments SF1 CS=4 g/min, P=450 bar, T=40 °C and SF5 CS=11 g/min, P=150 bar, T=40°C at two different times were selected. The profile of monosaccharides showed an important content of D-glucose (2.01±0.08 g/L) in comparison with the other monosaccharides such as D-galactose, D-mannose, D-arabinose, L-fucose and xylose Fig 5.

**Fig. 5.**
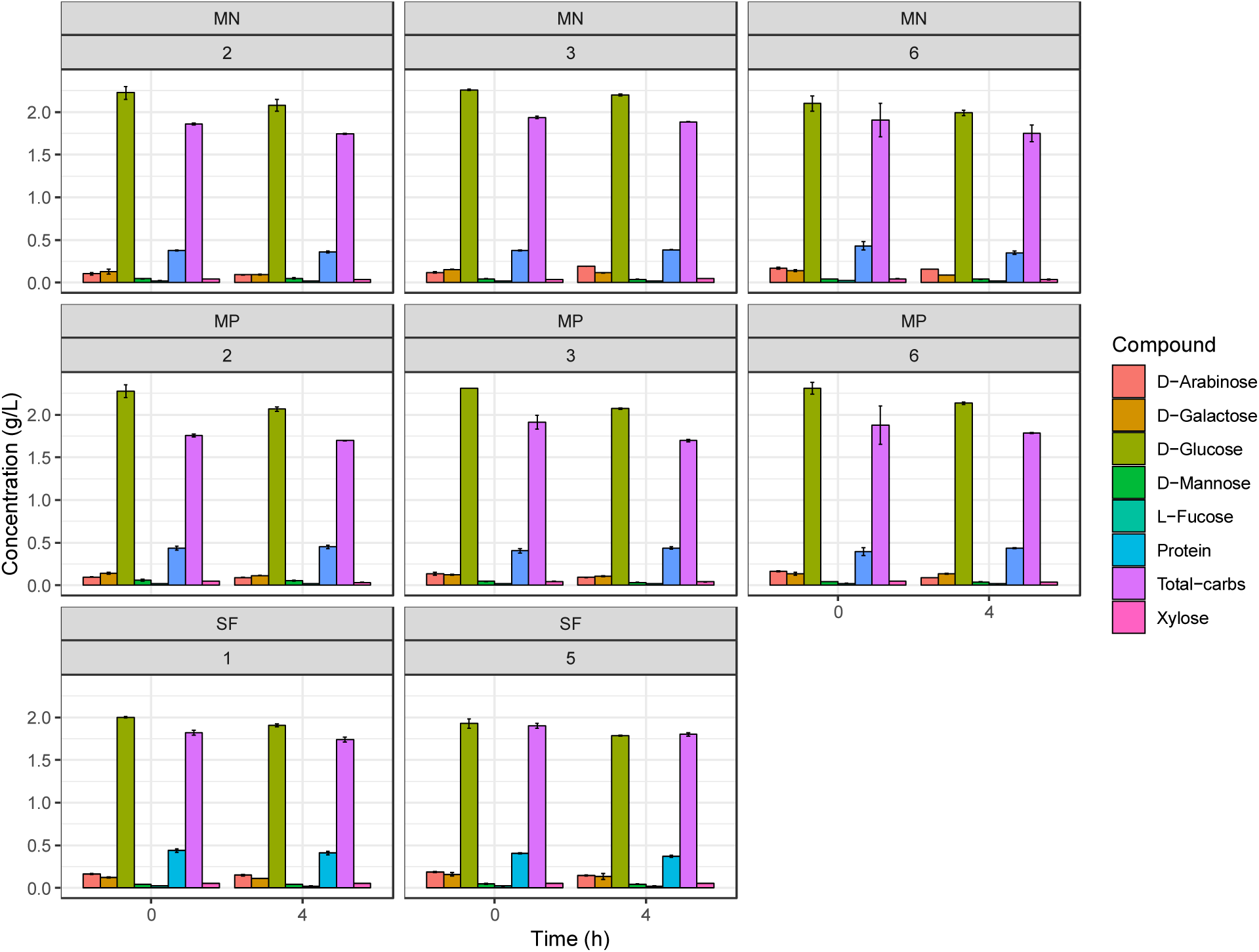
Total protein and monosaccharides concentration profile during fermentation of *Saccharomyces cerevisiae LPB-287* with cyanobacteria from SF, MN and MP pretreatments of selected samples. *(n=2).

Whereas, the MN experiments MN2 S=0.25, t=15 min, T=60 °C, MN3 S=0.25, t=55 min, T=40 °C and 6 S=0.81, t=15 min, T=60 °C at two different times were selected. The profile of monosaccharides shows an important content of D-glucose (2.48±0.06 g/L) in comparison with the other monosaccharides such as D-galactose, D-mannose, D-arabinose, L-fucose and xylose, Fig 5. However, in comparison with SF, MN has more content of D-glucose available for the fermentation process, due to the polarity of the solvents used in the extraction processes.

For MP, the experiments MP2 S=0.25, t=15 min, T=60 °C, 3 S=0.25, t=55 min, T=40 °C and MP6 S=0.81, t=15 min, T=60 °C at two different times were selected. The profile of monosaccharides shows an important content of D-glucose (2.31±0.04 g/L) in comparison with the other monosaccharides such as D-galactose, D-mannose, D-arabinose, L-fucose and xylose Fig 5. However, in comparison with SF and MN, MP has less content of D-glucose available for the fermentation process, and in terms of protein content, MP pretreatment showed a higher content than MN.

Furthermore, in all cases, the SF, MN and MP monosaccharides profile fits with the previous reports from the sugar content in *A. platensis* (Hahn et al., 2012; Shekharam et al., 1987).

*S. cerevisiae* ferments glucose, which represents the dominant sugar in almost all hydrolysates from biomass used as carbon source for ethanol production (Bakker et al., 2000). Also, mannose and fructose are two isomers of glucose present in the biomass hydrolysates that can be fermented by *S. cerevisiae* strains. As far as we know, the yeast capable of fermenting glucose and that can also ferment fructose and mannose, is known as the Kluyver rule (Van Dijken et al., 1986). Based on this rule, we assume that the strain *S. cerevisiae* LPB-287 can ferment mannose, but its consumption is mediated by the kinetics of mixed-substrate utilization because both compete for the same hexose transporters (Reifenberger et al., 1997). Furthermore, in our experiments we detected galactose Fig 5 but the glucose concentration causes a complete repression of galactose metabolism (Johnston et al., 1994), thus preventing the simultaneous consumption of glucose and galactose. However, sequential consumption of different monosaccharides can represent a problem in the ethanol production, due to the lag phase that separates glucose and galactose consumption in mixed-substrate cultures similar to our experiments. Therefore, this problem needs to be undertaken. Xylose and arabinose cannot be fermented or assimilated by *S. cerevisiae* LPB-87. Therefore, their concentration does not change along the experiments Fig 5.

Finally, concerning the protein content, no changes were observed among the selected samples (0.47±0.05 g/L), (0.38±0.05 g/L), (0.43±0.05 g/L) for SF, MN and MP Fig. 5, respectively.

### Effect of SFE and MAE pretreatment of cyanobacteria substrate on lactic acid production and consumption rate of reducing sugars

The depleted biomass was adopted as substrate for lactic acid fermentation by *L*. *acidophilus* ATCC 43121. As shown in Fig 6, the highest concentrations of lactic acid, 9.67±0.05 and 9.62±0.05 g/L, were achieved with the experiments SF8 and SF11, respectively.

**Fig. 6.**
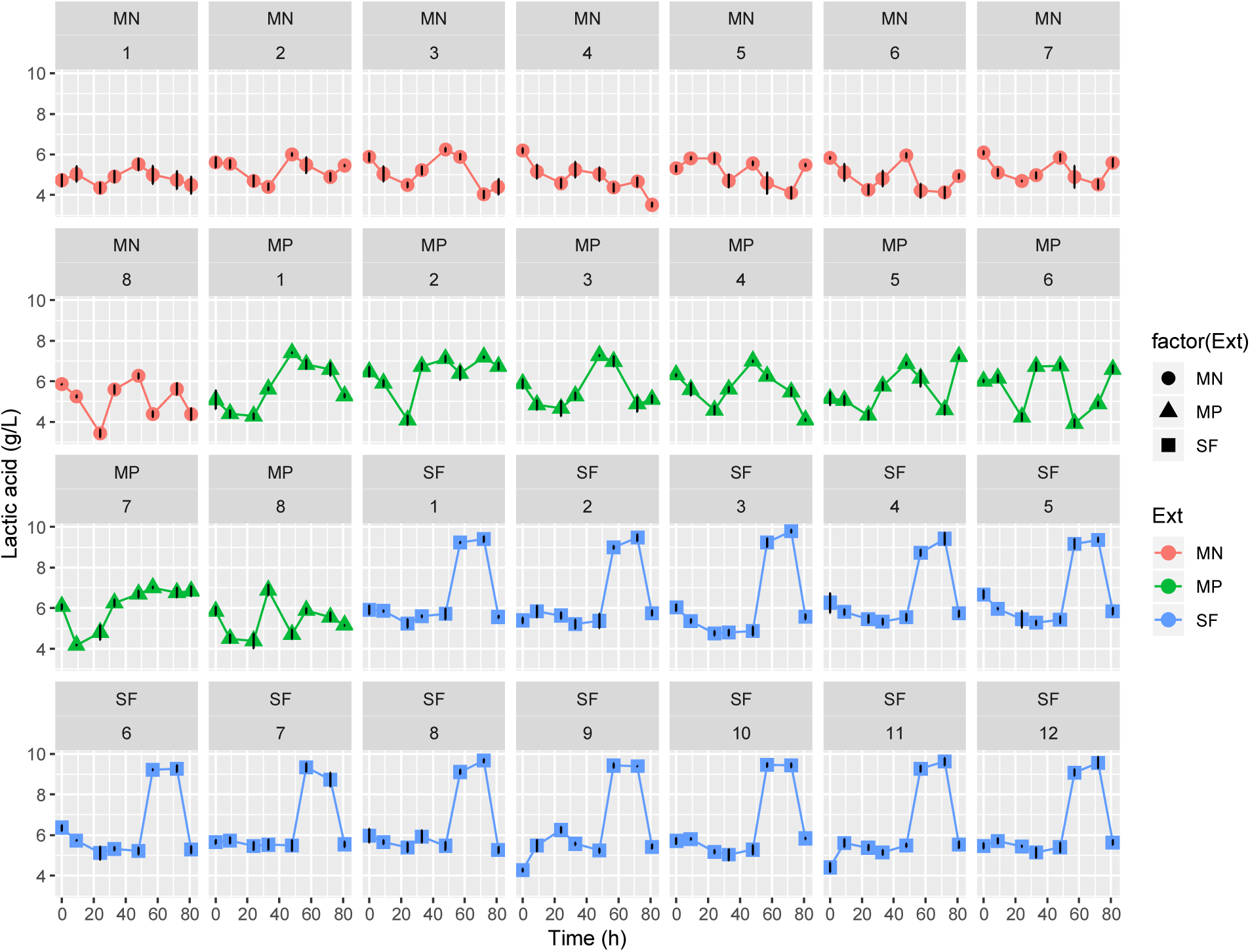
Time course of concentration and yield of lactic acid during fermentation of *Lactobacillus acidophilus* ATCC 43121 with cyanobacteria from SF, MN and MP pretreatments. *(factor(Ext) refers to the type of pretreatment respectively, n=3).

**Fig. 7.**
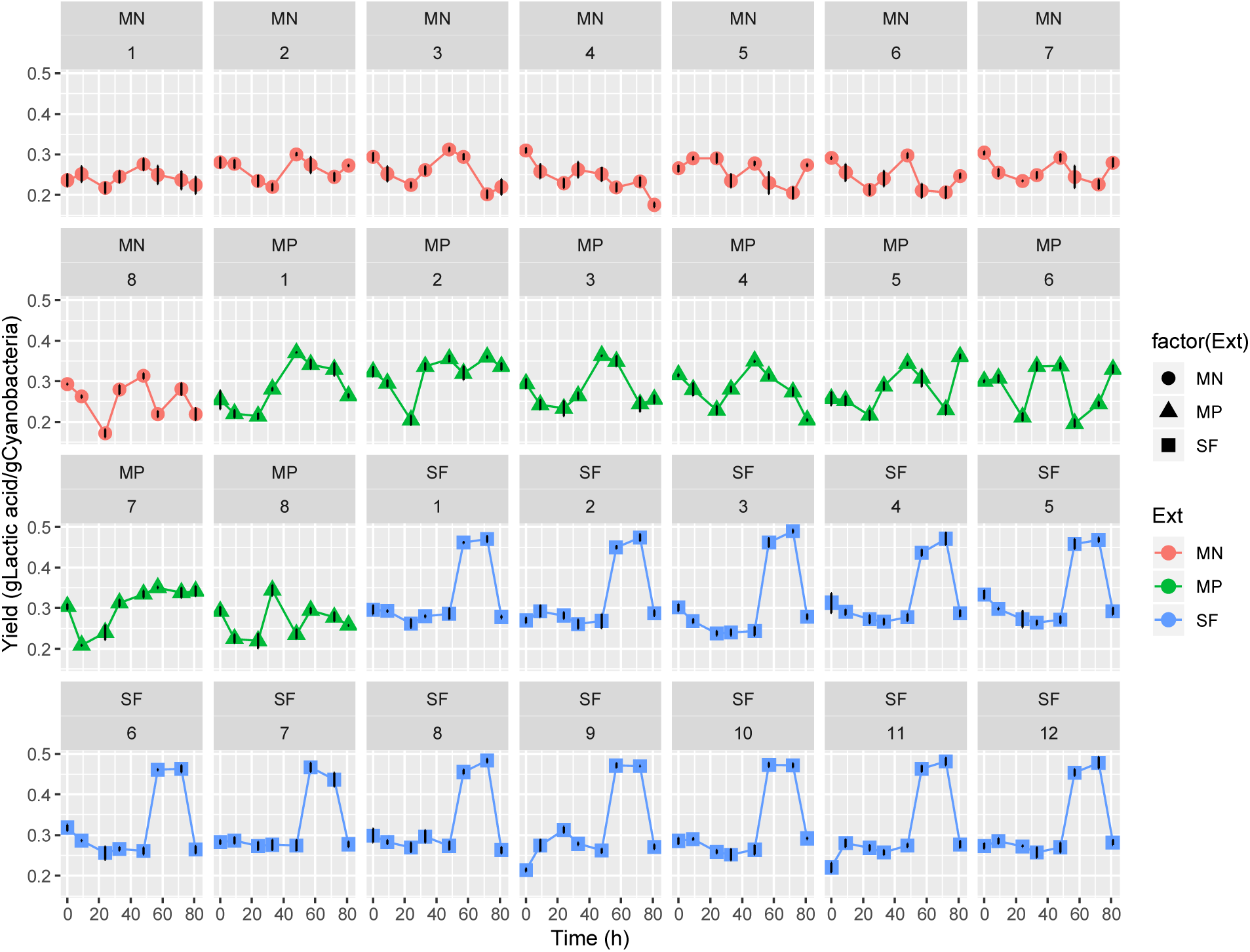
Time course of yield of lactic acid during fermentation of *Lactobacillus acidophilus* ATCC 43121 with depleted cyanobacteria biomass from SF, MN and MP pretreatments. *(factor(Ext) refers to the type of pretreatment respectively, n=3)

Regarding the lactic acid yield for SF, the highest yield (% gLactic acid/gCyanobacteria) was 0.48 in the experiments SF8 CS=11 g/min, P=450 bar, T=40 °C and 11 CS=11 g/min, P=150 bar, T=60 °C (Fig 7).

Whereas for MAE pretreatment, two sets of conditions, MN and MP, were evaluated as a substrate for lactic acid fermentation by *L*. *acidophilus* ATCC 43121 after the pretreatment with MAE. According to Fig 6, for MN set, the highest lactic acid concentration was obtained in the experiment MN3, S=0.25, t=55 min, T=40 °C (6.24±0.08 g/L). Regarding the lactic acid yield, the highest value (% gLactic acid/gCyanobacteria) of 0.31 was obtained in the experiment MN3 S=0.25, t=55 min, T=40 °C, Fig. 7.

Concerning the MP experiments, the highest lactic acid concentration was obtained in the experiment MP3 S=0.25, t=55 min, T=40 °C, (7.26±0.05 g/L), Fig 6. Regarding the lactic acid yield for MP, the highest value (% gLactic acid/gCyanobacteria) of 0.37 was obtained in the experiment MP3 S=0.25, t=55 min, T=40 °C, Fig 7, Table 1 (b).

To the extent of our knowledge, there are not studies of lactic production from *A. platensis* depleted biomass. However, in this frame, exist some studies based on other algal cultures. It has been reported the use of lyophilized biomass of the *A. platensis* as the substrate for lactic acid fermentation by the probiotic bacterium *Lactobacillus plantarum* ATCC 8014 with a lactic acid concentration of 3.7 g L^−1^ (Niccolai et al., 2019). Moreover, algal carcass (AC) a low-value byproduct of algae after its conversion to biodiesel was used to ferment algal carcass media (ACM), including 2% ACM with 1.9% glucose and 2 g/L amino acid mixture (ACM-GA) with *Lactobacillus delbrueckii* ssp. bulgaricus ATCC 11842. The results showed that the highest content of lactic acid was 3.31 g/L (Li et al., 2016). Furthermore, biomass from *Enteromorpha prolifera* was hydrolyzed with 0.5 M sulfuric acid at 120°C for 2h, and therefore was used as carbon source for a fermentation process with *Lactobacillus rhamnosus*, the results showed that the highest content of lactic acid was 4.3 g/L (Hwang et al., 2012). Also, biomass from *Gelidium amansii* was hydrolyzed with 3% v/v sulfuric acid at 140°C for 5 min, therefore the hydrolysate was fermented with *L. rhamnosus* and the highest content of lactic acid was 12.5 g/L (Jang et al., 2013).

In reference to this information, we have a higher content of lactic acid than the results from other algal cultures such as algal carcass and *Enteromorpha prolifera,* but we have a lower content in comparison with the use of *Gelidum amansii* biomass. These results can be due to the different composition of algal biomass, since they are part of a big biological group with several differences between them.

Additionally, according to our previous reports (Esquivel-Hernández et al., 2017a), similarly to the ethanol results, the SF experiment SF8 CS=11 g/min, P=450 bar, Table 1 (a) offers the best option in terms of suitability for the integrated biorefinery due to the highest extraction yield (7.48 % w/w). While for MN, the lactic acid content is lower than the results from SF pretreatment due to the differences between the extraction processes in terms of the types of solvents (Supercritical carbon dioxide vs limonene, ethyl acetate, hexane i.e.). According to our previous reports (Esquivel-Hernández et al., 2017b), experiment MN3 S=0.25, t=55 min, T=40 °C represents a fair option for the integrated biorefinery due to its good extraction yield (4.67 % w/w) of bioactive compounds and its specific conditions Table 1 (b), while the contents of lactic acid in MP pretreatments were lower than the SF and MN values.

Regarding the assimilation of cyanobacterial biomass by *L. acidophilus* ATCC 43121, we measured the change in reducing sugars content in the time. These results are presented in Fig 8., for SFE all extraction samples evaluated showed a decrease in glucose content when compared with control biomass Fig 8. This behavior can be due to metabolism of the genus *Lactobacillus* since the hexose sugars are rapidly metabolized by *Lactobacillus*; however, the depleted cyanobacterial biomass contains other kind of sugars such as pentose sugars that *L. acidophilus* does not use when there are the hexoses sugars. Hexose sugars are considered to be efficiently fermented to lactic acid, whereas pentose sugars cannot be consumed (Abdel-Rahman et al., 2011). The highest content of reducing sugars was obtained under the experiments SF5 and SF10 Table 1 (a), Fig 8.

**Fig. 8.**
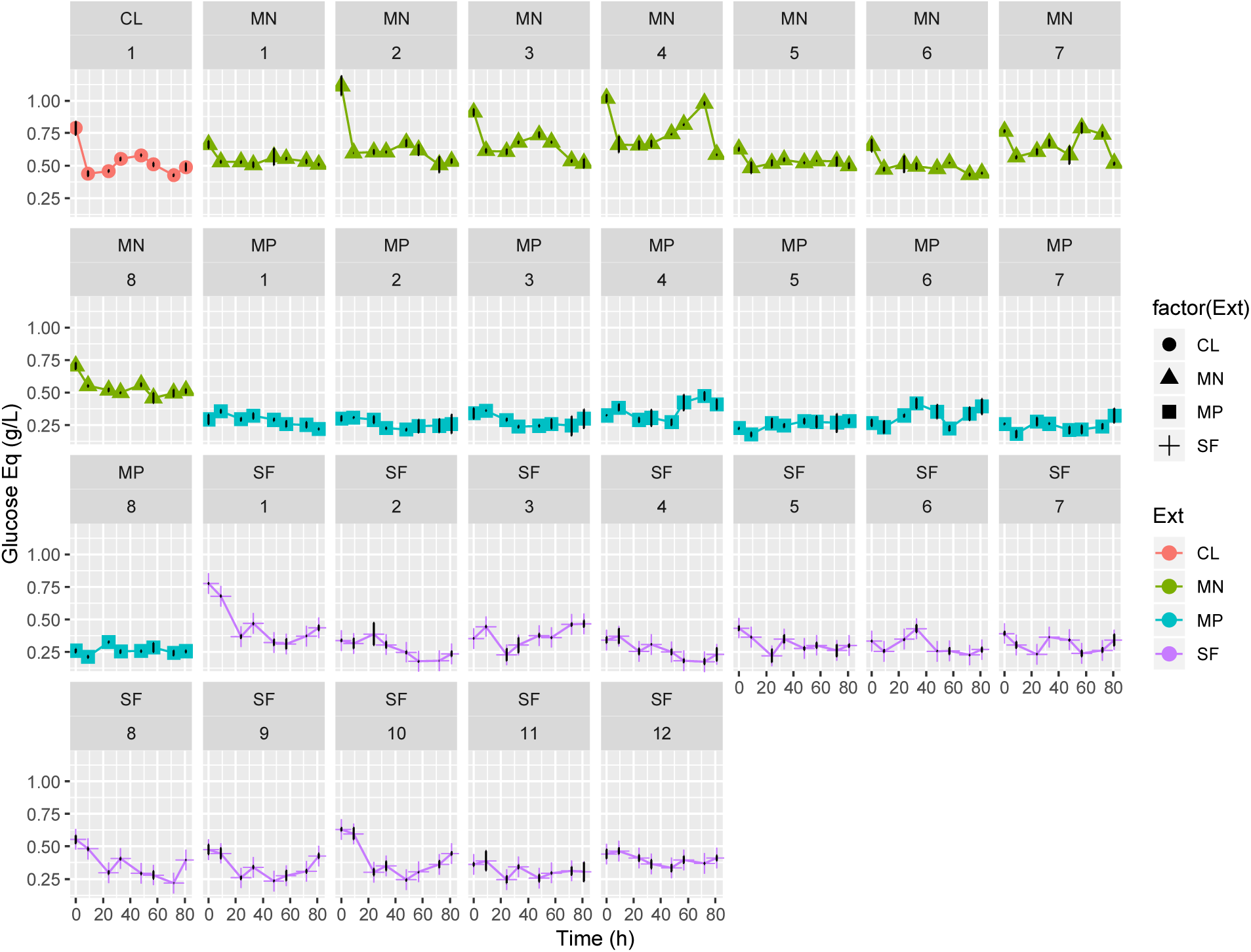
Time course of reducing sugar concentration measured by DNS method during fermentation of *Lactobacillus acidophilus* ATCC 43121 with cyanobacteria from SF, MN and MP pretreatments. *(factor(Ext) refers to the type of pretreatment, respectively, n=3, CL is a positive control).

The results of MN depleted biomass assimilation by *L. acidophilus* ATCC 43121 are showed in Fig 8. All these samples obtained by MN show a remarkable increase in reducing sugars when compared with control biomass. This can be due to the extraction mixture used for MN, which comprises limonene and ethyl acetate (Esquivel-Hernández et al., 2017b) which are non-polar solvents and as a result of their polarity, we only extract non polar compounds, therefore the concentration of carbohydrates in the biomass increases. Differently to pretreatment with SF and MP processes, in these results we can observe a major depletion in the reducing sugars after the first 10 hours. In this case, we can assume that hexose sugars are rapidly metabolized (Abdel-Rahman et al., 2011). Also, it is important to distinguish that these samples exhibited a different behavior of the control sample, mainly because in this pretreatment we started at the higher level of reducing sugars. The maximum content of reducing sugars was obtained under the experiments MN2 S=0.25, t=15 min, T=60 °C and MN3 S=0.25, t=55 min, T=40 °C Table 1(b).

The assimilation of MP depleted biomass by *L. acidophilus* ATCC 43121 measured as a change in reducing sugars are shown in Fig 8. As it is possible to see, all extraction samples obtained by MP show a remarkable decrease in reducing sugars when compared with control biomass. This can be due to the extraction mixture used for MP, which comprises water (10 mM ammonium acetate) and ethanol. Differently from pretreatment with SF processes, in these results we can observe a slight increase in the reducing sugars after the first 24 hours. However, after these 24 hours, we found a decrease in the reduced sugar contents. this can be due to a reduction in the availability of hexose sugars (Abdel-Rahman et al., 2011). The highest content of reducing sugars was obtained under the experiments MP3 S=0.25, t=55 min, T=40 °C and MP4 S=0.25, t=55 min, T=60 °C Table 1 (b). Differently from the ethanol results, the biomass that results from MP process represents a fair option for the lactic acid production in the frame of integrated biorefinery due to their higher content of lactic acid and the diversity of high value metabolites content (Esquivel-Hernández et al., 2017b). However, in the broad view, the SF pretreatment gives the highest content of lactic acid.

Our study demonstrates that the biomass resulting from this extraction process has the advantage of being appropriate as carbon source for lactic acid production because some of its carbohydrates have been released and are readily available for the *Lactobacillus* to ferment.

### Time of course of monosaccharides and protein, in selected conditions of lactic acid fermentation

The time of course of monosaccharides and protein of the most representative samples of SF, MN and MP pretreatments in the fermentation by *L. acidophilus* was performed. For SF, the experiments SF1 CS=4 g/min, P=450 bar, T=40 °C and SF5 CS=11 g/min, P=150 bar, T=60 °C at two different times were selected. The profile of monosaccharides shows an important content of D-glucose (1.46±0.07 g/L) in comparison with the other monosaccharides such as D-galactose, D-mannose, D-arabinose, L-fucose and xylose Fig 9.

**Fig. 9.**
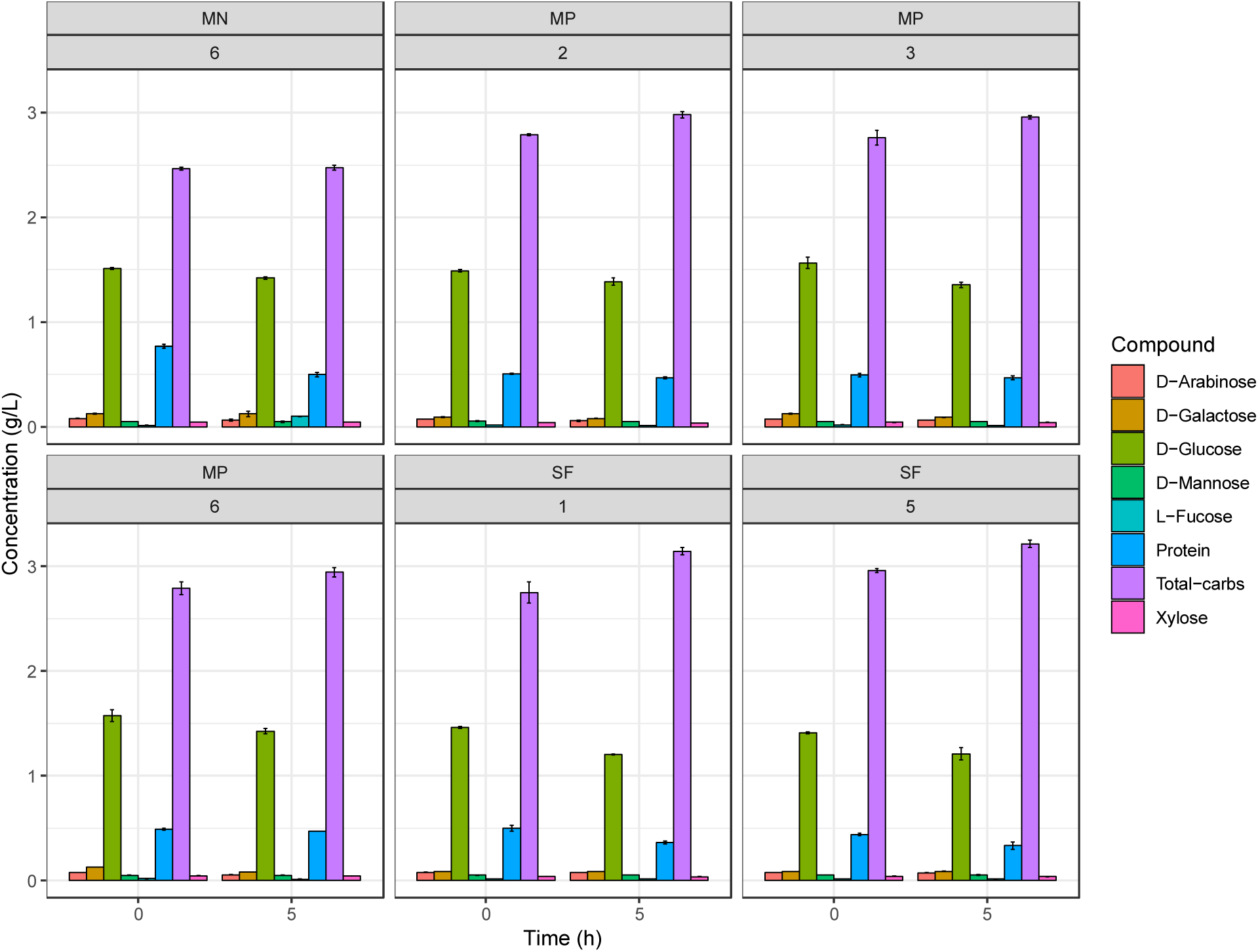
Total protein and monosaccharides concentration profile during fermentation of *Lactobacillus acidophilus* ATCC 43121 with cyanobacteria from SF, MN and MP pretreatments of selected samples. *(n=2).

For MN, the experiment MN6 S=0.81, t=15 min, T=60 °C at two different times was selected. The profile of monosaccharides shows an important content of D-glucose (1.63±0.07 g/L) in comparison with the other monosaccharides such as D-galactose, D-mannose, D-arabinose, L-fucose and xylose Fig 9.

For MP, the experiments MP2 S=0.25, t=15 min, T=60 °C, MP3 S=0.25, t=55 min, T=40 °C and MP6 S=0.81, t=15 min, T=60 °C at two different times were selected. The profile of monosaccharides shows an important content of D-glucose (1.57±0.06 g/L) in comparison with the other monosaccharides such as D-galactose, D-mannose, D-arabinose, L-fucose and xylose Fig 9.

Moreover, in all cases, SF, MN and MP fits with the previous reports from the sugar content in *A. platensis* (González-Fernández & Ballesteros, 2012). However, in comparison with SF, MN and MP have more content of D-glucose available for the fermentation process.

*L. acidophilus* is an obligated homo-fermenter and is among the least oxygen-tolerant lactobacilli (Claesson et al., 2007). Also, it has been identified and characterized different transporters that allows the uptake of several carbohydrates such as glucose, fructose and trehalose, and this catabolic machinery is highly regulated at the transcription level, suggesting that the *L. acidophilus* transcriptome is flexible, dynamic and designed for efficient carbohydrate utilization. Furthermore *L. acidophilus* appears able to adapt its metabolic machinery to fluctuation carbohydrate sources available in the environment (Barrangou et al., 2006). Based on this information, *L. acidophilus* represent a suitable microorganism for the production of lactic acid in the frame of integrated biorefinery with cyanobacterial biomass.

In terms of the protein content, no changes were observed among the selected samples for SF (0.44±0.06 g/L), and MP (0.50±0.04 g/L), but in the samples from MN we observed a reduction in the content of total protein (Fig 9).

## Conclusion

Ethanol and lactic acid possess high market value. The first one can be used as a biofuel and the former possess a broader range of applications in the food and chemical industries. The main drawback for the bulk production of these molecules are the carbon source and the cost of the elements present in their culture media. To overcome these drawbacks, an integrated biorefinery of the production of high value metabolites can accelerate their production at industrial level.

The cyanobacterial biomass has proven to resolve the problems that are being encountered with the current fermentation feedstocks through the provision of several advantages and cost benefits. The present study allowed demonstrating that the biomass resulting from SFE or MAE process has the advantage of being used as carbon source for ethanol and lactic acid production because some of its carbohydrates have been released and are readily available for the fermentation processes. For ethanol, MN pretreatment showed the highest content (3.02 ±0.07 g/L), while for lactic acid, SF pretreatment showed the highest content (9.67±0.05). The application of the green extraction process to produce high value metabolites and the subsequent use of the depleted biomass has been confirmed the suitability for the development of an integrated biorefinery with *A. platensis* as a feedstock. To the best of our knowledge, this is the first study in which *A. platensis* biomass depleted by SFE and MAE treatments are used as feedstock for bioethanol and lactic acid production.

## Acknowledgments

This research was supported by a Marie Curie International Research Staff Exchange Scheme Fellowship within the 7th European Community Framework Programme: “Improvement of technologies and tools, e.g. biosystems and biocatalysts, for waste conversion to develop an assortment of high added value eco-friendly and cost-effective bio-products” BIOASSORT (contract number 318931). *S. cerevisiae LPB-287* and *L. acidophilus* ATCC 43121 were kindly provided by Carlos Ricardo Soccol and Luciana Porto de Souza Vandenberghe from the Federal University of Parana, Brazil. The authors are grateful to Rossana Liguori, Gabriela Cerullo, Davide Agostino Cecchini, and Ginevra del Vecchio for technical assistance with the experimental setup.

## References

Abdel-Rahman, M.A., Tashiro, Y., Sonomoto, K. 2011. Lactic acid production from lignocellulose-derived sugars using lactic acid bacteria: overview and limits. Journal of biotechnology, 156(4), 286–301.

Aikawa, S., Joseph, A., Yamada, R., Izumi, Y., Yamagishi, T., Matsuda, F., Kawai, H., Chang, J.-S., Hasunuma, T., Kondo, A. 2013. Direct conversion of Spirulina to ethanol without pretreatment or enzymatic hydrolysis processes. Energy & Environmental Science, 6(6), 1844–1849.

Bakker, B.M., Bro, C., Kötter, P., Luttik, M.A., Van Dijken, J.P., Pronk, J.T. 2000. The mitochondrial alcohol dehydrogenase Adh3p is involved in a redox shuttle in Saccharomyces cerevisiae. Journal of bacteriology, 182(17), 4730–4737.

Barrangou, R., Azcarate-Peril, M.A., Duong, T., Conners, S.B., Kelly, R.M., Klaenhammer, T.R. 2006. Global analysis of carbohydrate utilization by Lactobacillus acidophilus using cDNA microarrays. Proceedings of the National Academy of Sciences of the United States of America, 103(10), 3816–3821.

Campanella, L., Crescentini, G., Avino, P. 1999. Chemical composition and nutritional evaluation of some natural and commercial food products based on Spirulina. Analusis, 27(6), 533–540.

Capolupo, L., Faraco, V. 2016. Green methods of lignocellulose pretreatment for biorefinery development. Applied microbiology and biotechnology, 100(22), 9451–9467.

Claesson, M.J., Van Sinderen, D., O’Toole, P.W. 2007. The genus Lactobacillus–a genomic basis for understanding its diversity. FEMS microbiology letters, 269(1), 22–28.

Dixit, R.B., Suseela, M. 2013. Cyanobacteria: Potential candidates for drug discovery. Antonie Van Leeuwenhoek, 103(5), 947–961.

Ediriweera, M.K., Tennekoon, K.H., Samarakoon, S.R. 2017. A Review on Ethnopharmacological Applications, Pharmacological Activities, and Bioactive Compounds of Mangifera indica (Mango). Evidence-Based Complementary and Alternative Medicine, 2017.

Esquivel-Hernández, D.A., López, V.H., Rodríguez-Rodríguez, J., Alemán-Nava, G.S., Cuéllar-Bermúdez, S.P., Rostro-Alanis, M., Parra-Saldívar, R. 2016. Supercritical carbon dioxide and microwave-assisted extraction of functional lipophilic compounds from Arthrospira platensis. International journal of molecular sciences, 17(5), 658.

Esquivel-Hernández, D.A., Rodríguez-Rodríguez, J., Cuéllar-Bermúdez, S.P., García-Pérez, J.S., Mancera-Andrade, E.I., Núñez-Echevarría, J.E., Ontiveros-Valencia, A., Rostro-Alanis, M., García-García, R.M., Torres, J.A. 2017a. Effect of Supercritical Carbon Dioxide Extraction Parameters on the Biological Activities and Metabolites Present in Extracts from Arthrospira platensis. Marine drugs, 15(6), 174.

Esquivel-Hernández, D.A., Rodríguez-Rodríguez, J., Rostro-Alanis, M., Cuéllar-Bermúdez, S.P., Mancera-Andrade, E.I., Núñez-Echevarría, J.E., García-Pérez, J.S., Chandra, R., Parra-Saldívar, R. 2017b. Advancement of green process through microwave-assisted extraction of bioactive metabolites from Arthrospira Platensis and bioactivity evaluation. Bioresource technology, 224, 618–629.

Esquivel-Hernández, D.A., Ibarra-Garza, I.P., Rodríguez-Rodríguez, J., Cuéllar-Bermúdez, S.P., Rostro-Alanis, M.d.J., Alemán-Nava, G.S., García-Pérez, J.S., Parra-Saldívar, R. 2017. Green extraction technologies for high-value metabolites from algae: a review. Biofuels, Bioproducts and Biorefining, 11(1), 215–231.

González-Fernández, C., Ballesteros, M. 2012. Linking microalgae and cyanobacteria culture conditions and key-enzymes for carbohydrate accumulation. Biotechnology advances, 30(6), 1655–1661.

Hahn, T., Lang, S., Ulber, R., Muffler, K. 2012. Novel procedures for the extraction of fucoidan from brown algae. Process biochemistry, 47(12), 1691–1698.

Harun, R., Danquah, M.K., Forde, G.M. 2010. Microalgal biomass as a fermentation feedstock for bioethanol production. Journal of chemical technology and biotechnology, 85(2), 199–203.

Hossain, M.N.B., Basu, J.K., Mamun, M. 2015. The production of ethanol from microalgae Spirulina. Procedia Engineering, 105, 733–738.

Hwang, H.J., Kim, S.M., Chang, J.H., Lee, S.B. 2012. Lactic acid production from seaweed hydrolysate of Enteromorpha prolifera (Chlorophyta). Journal of applied phycology, 24(4), 935–940.

Jang, S.-S., Shirai, Y., Uchida, M., Wakisaka, M. 2013. Potential use of Gelidium amansii acid hydrolysate for lactic acid production by Lactobacillus rhamnosus. Food Technology and Biotechnology, 51(1), 131.

Johnston, M., Flick, J.S., Pexton, T. 1994. Multiple mechanisms provide rapid and stringent glucose repression of GAL gene expression in Saccharomyces cerevisiae. Molecular and cellular biology, 14(6), 3834–3841.

Kim, H.M., Oh, C.H., Bae, H.-J. 2017. Comparison of red microalgae (Porphyridium cruentum) culture conditions for bioethanol production. Bioresource technology, 233, 44–50.

Kim, J.-H., Block, D.E., Mills, D.A. 2010. Simultaneous consumption of pentose and hexose sugars: an optimal microbial phenotype for efficient fermentation of lignocellulosic biomass. Applied microbiology and biotechnology, 88(5), 1077–1085.

Kumari, D.J., Babitha, B., Jaffar, S., Prasad, M.G., Ibrahim, M., Khan, M. 2011. Potential health benefits of Spirulina platensis. Int. J. Adv. Pharm. Sci, 2, 417–422.

Li, C., Zhang, G., Mao, X., Wang, J., Duan, C., Wang, Z., Liu, L. 2016. Growth and acid production of Lactobacillus delbrueckii ssp. bulgaricus ATCC 11842 in the fermentation of algal carcass. Journal of dairy science, 99(6), 4243–4250.

Liguori, R., Amore, A., Faraco, V. 2013. Waste valorization by biotechnological conversion into added value products. Applied microbiology and biotechnology, 97(14), 6129–6147.

Liguori, R., Faraco, V. 2016. Biological processes for advancing lignocellulosic waste biorefinery by advocating circular economy. Bioresource technology, 215, 13–20.

Liguori, R., Soccol, C.R., Porto de Souza Vandenberghe, L., Woiciechowski, A.L., Faraco, V. 2015a. Second generation ethanol production from brewers’ spent grain. Energies, 8(4), 2575–2586.

Liguori, R., Soccol, C.R., Vandenberghe, L.P.d.S., Woiciechowski, A.L., Ionata, E., Marcolongo, L., Faraco, V. 2015b. Selection of the strain Lactobacillus acidophilus ATCC 43121 and its application to brewers’ spent grain conversion into lactic acid. BioMed research international, 2015.

Markou, G., Angelidaki, I., Nerantzis, E., Georgakakis, D. 2013. Bioethanol production by carbohydrate-enriched biomass of Arthrospira (Spirulina) platensis. Energies, 6(8), 3937–3950.

Miller, G.L. 1959. Use of dinitrosalicylic acid reagent for determination of reducing sugar. Analytical chemistry, 31(3), 426–428.

Mondal, M., Goswami, S., Ghosh, A., Oinam, G., Tiwari, O., Das, P., Gayen, K., Mandal, M., Halder, G. 2017. Production of biodiesel from microalgae through biological carbon capture: a review. 3 Biotech, 7(2), 99.

Ngamsirisomsakul, M., Reungsang, A., Liao, Q., Kongkeitkajorn, M.B. 2019. Enhanced bio-ethanol production from Chlorella sp. biomass by hydrothermal pretreatment and enzymatic hydrolysis. Renewable Energy, 141, 482–492.

Nguyen, C.M., Kim, J.-S., Song, J.K., Choi, G.J., Choi, Y.H., Jang, K.S., Kim, J.-C. 2012. D-Lactic acid production from dry biomass of Hydrodictyon reticulatum by simultaneous saccharification and co-fermentation using Lactobacillus coryniformis subsp. torquens. Biotechnology letters, 34(12), 2235–2240.

Niccolai, A., Shannon, E., Abu-Ghannam, N., Biondi, N., Rodolfi, L., Tredici, M.R. 2019. Lactic acid fermentation of Arthrospira platensis (spirulina) biomass for probiotic-based products. Journal of Applied Phycology, 31(2), 1077–1083.

Reifenberger, E., Boles, E., Ciriacy, M. 1997. Kinetic characterization of individual hexose transporters of Saccharomyces cerevisiae and their relation to the triggering mechanisms of glucose repression. The FEBS Journal, 245(2), 324–333.

Shekharam, K.M., Venkataraman, L., Salimath, P. 1987. Carbohydrate composition and characterization of two unusual sugars from the blue green alga Spirulina platensis. Phytochemistry, 26(8), 2267–2269.

Shokrkar, H., Ebrahimi, S., Zamani, M. 2017. Bioethanol production from acidic and enzymatic hydrolysates of mixed microalgae culture. Fuel, 200, 380–386.

Singh, R., Parihar, P., Singh, M., Bajguz, A., Kumar, J., Singh, S., Singh, V.P., Prasad, S.M. 2017. Uncovering potential applications of cyanobacteria and algal metabolites in biology, agriculture and medicine: current status and future prospects. Frontiers in microbiology, 8, 515.

Sosa-Hernández, J.E., Romero-Castillo, K.D., Parra-Arroyo, L., Aguilar-Aguila-Isaías, M.A., García-Reyes, I.E., Ahmed, I., Parra-Saldivar, R., Bilal, M., Iqbal, H. 2019. Mexican Microalgae Biodiversity and State-Of-The-Art Extraction Strategies to Meet Sustainable Circular Economy Challenges: High-Value Compounds and Their Applied Perspectives. Marine drugs, 17(3), 174.

Van Dijken, J.P., Van Den Bosch, E., Hermans, J.J., De Miranda, L.R., Scheffers, W.A. 1986. Alcoholic fermentation by ‘non-fermentative’yeasts. Yeast, 2(2), 123–127.

Van Eykelenburg, C. 1977. On the morphology and ultrastructure of the cell wall of Spirulina platensis. Antonie van leeuwenhoek, 43(2), 89–99.

Wang, F., Ma, Y., Liu, Y., Cui, Z., Ying, X., Zhang, F., Linhardt, R.J. 2017. A simple strategy for the separation and purification of water-soluble polysaccharides from the fresh Spirulina platensis. Separation Science and Technology, 52(3), 456–466.

